# D614G Substitution of SARS-CoV-2 Spike Protein Increases Syncytium Formation and Viral Transmission via Enhanced Furin-mediated Spike Cleavage

**DOI:** 10.1101/2021.01.27.428541

**Authors:** Ya-Wen Cheng, Tai-Ling Chao, Chiao-Ling Li, Sheng-Han Wang, Han-Chieh Kao, Ya-Min Tsai, Hurng-Yi Wang, Chi-Ling Hsieh, Pei-Jer Chen, Sui-Yuan Chang, Shiou-Hwei Yeh

**Author notes:** These authors contributed equally (Ya-Wen Cheng, Tai-Ling Chao). Corresponding author: S.Y.C.; S.H.Y., **CORRESPONDENCE:** Sui-Yuan Chang, Ph.D., Professor, Department of Laboratory Medicine, National Taiwan University College of Medicine, No. 7 Chung-Shan South Road, Taipei 100, Taiwan. or Shiou-Hwei Yeh, Ph.D., Professor, Department of Microbiology, National Taiwan University College of Medicine, No. 1, Jen-Ai Road, Section 1, Taipei 100, Taiwan.

## Abstract

Since the D614G substitution in the spike (S) of SARS-CoV-2 emerged, the variant strain underwent rapid expansion to become the most abundant strain worldwide. Therefore, this substitution may provide an advantage of viral spreading. To explore the mechanism, we analyzed 18 viral isolates containing S proteins with either G614 or D614. Both the virus titer and syncytial phenotype were significantly increased in S-G614 than in S-D614 isolates. We further showed increased cleavage of S at the furin substrate site, a key event that promotes syncytium, in S-G614 isolates. These functions of the D614G substitution were validated in cells expressing S protein. The effect on syncytium was abolished by furin inhibitor treatment and mutation of the furin-cleavage site, suggesting its dependence on cleavage by furin. Our study provides a mechanistic explanation for the increased transmissibility of S-G614 containing SARS-CoV-2 through enhanced furin-mediated S cleavage, which increases membrane fusion and virus infectivity.

## INTRODUCTION

The severe acute respiratory syndrome coronavirus 2 (SARS-CoV-2) infection causes a rapid accumulation of confirmed and fatal cases and poses a threat to public health around the world. A better understanding of viral evolution and characterization of viral genetic variations usually provides valuable insights into the mechanisms linked to pathogenesis, antiviral drug resistance, and immune responses, which also impact the development of new vaccines, antiviral drugs and diagnostic tests (Chellapandi and Saranya, 2020; Sanjuan and Domingo-Calap, 2016; Young et al., 2020). Therefore, analysis of viral genomes and monitoring of the evolutionary trajectory of SARS-CoV-2 over time have been meticulously conducted to identify any specific genetic variations that contribute to the transmissibility and virulence of SARS-CoV-2 (Chitranshi et al., 2020; Mercatelli and Giorgi, 2020; Pachetti et al., 2020).

Among the genetic variations that have evolved during the course of the SARS-CoV-2 outbreak, the D614G substitution in the spike (S) protein, which corresponds to a change of the A nucleotide at genome position 23,403 to a G, has been identified as the signature of the A2a clade of SARS-CoV-2, the most prevalent clade (Korber et al., 2020; Yin, 2020). This substitution emerged at low frequency in early March but had rapidly expanded to become the most abundant clade worldwide by April to May (Korber et al., 2020), which was thus proposed to provide a selective fitness advantage during the outbreak. Korber B *et al.* recently reported that the S-G614-containing strain is associated with a higher viral load in the upper respiratory tract in infected individuals, though not increased disease severity (Korber et al., 2020). Several recent reports further demonstrated that pseudotyped viruses or the engineered viruses containing the S-G614 exhibit significantly higher infectivity than those containing S-D614 (Daniloski et al., 2020; Hu et al., 2020; Plante et al., 2020; Weissman et al., 2020). Sera from most convalescent COVID-19 patients could neutralize both S-D614 and S-G614 pseudotyped viruses with comparable efficiencies (Korber et al., 2020). These findings thus raised the possibility that D614G substitution confers increased infectivity and transmissibility, promoting its rapid expansion worldwide, but not through an increased binding affinity for ACE2 or increased escape of immune surveillance. Elucidating the molecular basis for the higher infectivity of D614G virus is urgent to understand its predominance and design an effective treatment strategy for patients.

As documented, release of the fusion peptide from the S protein via cleavage by host proteases, including furin/proprotein convertases (PCs) at the S1/S2 boundary and TMPRSS2 at the S2’ site within the S2 domain, is a prerequisite for the membrane fusion of SARS-CoV-2 with the target cells and thus viral infection (Hoffmann et al., 2020; Hou et al., 2020b; Ou et al., 2016). In our recent study, we found that the cleavage of S by furin/PCs at the S1/S2 boundary is also critical for S-mediated syncytium formation, another pathogenic event that contributes to increased viral transmission (Cheng et al., 2020). Meanwhile, a recent cryogenic electron microscopy (cryo-EM) analysis suggested that the D614G substitution induces a conformational change in the S protein (Yurkovetskiy et al., 2020). As noted, the D614G substitution is located at the C-terminal region of the S1 domain of the S protein, close to the furin-cleavage site (between a.a. 685-686). It thus raised a possibility that the D614G substitution might contribute to increase accessibility of the S protein for cleavage by furin through a conformational change and thus increasing the membrane fusion activity, as the basis for the increased infectivity and transmission capability of S-G614-containing SARS-CoV-2. To test this hypothesis, we first compared the virus titer, syncytial phenotype and cleavage of spike protein in 18 clinical SARS-CoV-2 isolates. The effects of the D614G substitution on enhanced syncytium and spike cleavage have been further quantitatively validated in the cells expressing spike protein. This *in vitro* assay system has been further used to examine the critical role of furin mediated S cleavage for the effect of D614G substitution. We expect the findings will provide a mechanistic explanation for the rapid expansion of S-G614 containing SARS-CoV-2 and help develop therapeutic strategy for intervening their spreading in population.

## RESULTS

### The virus titer and syncytial phenotype were significantly higher in S-G614 containing viral isolates than in S-D614 containing viral isolates

We first compared the virus titer and syncytial phenotype of 18 clinical SARS-CoV-2 isolates containing either S-D614 or S-G614 protein from infected Vero E6 cells (NTU01 to NUT18, Table S1). Interestingly, we found that the S-G614-containing viruses (n=10) had a significantly higher virus titer than the S-D614-containing viruses (n=8), as determined by plaque assay (Figure 1A). Consistently, northern blotting and qRT-PCR analysis confirmed the higher viral RNA levels in Vero E6 cells infected with S-G614-containing viruses compared to S-D614-containing viruses (*P*=0.0164) (Figure 1B). Microscopic observation further showed a higher syncytium level in S-G614-containing viruses than in S-D614-containing viruses (Figure 1C). These results obtained with the virus isolates suggest the possible functional effect of the D614G substitution in increasing virus production and syncytium formation.

**Figure 1.**
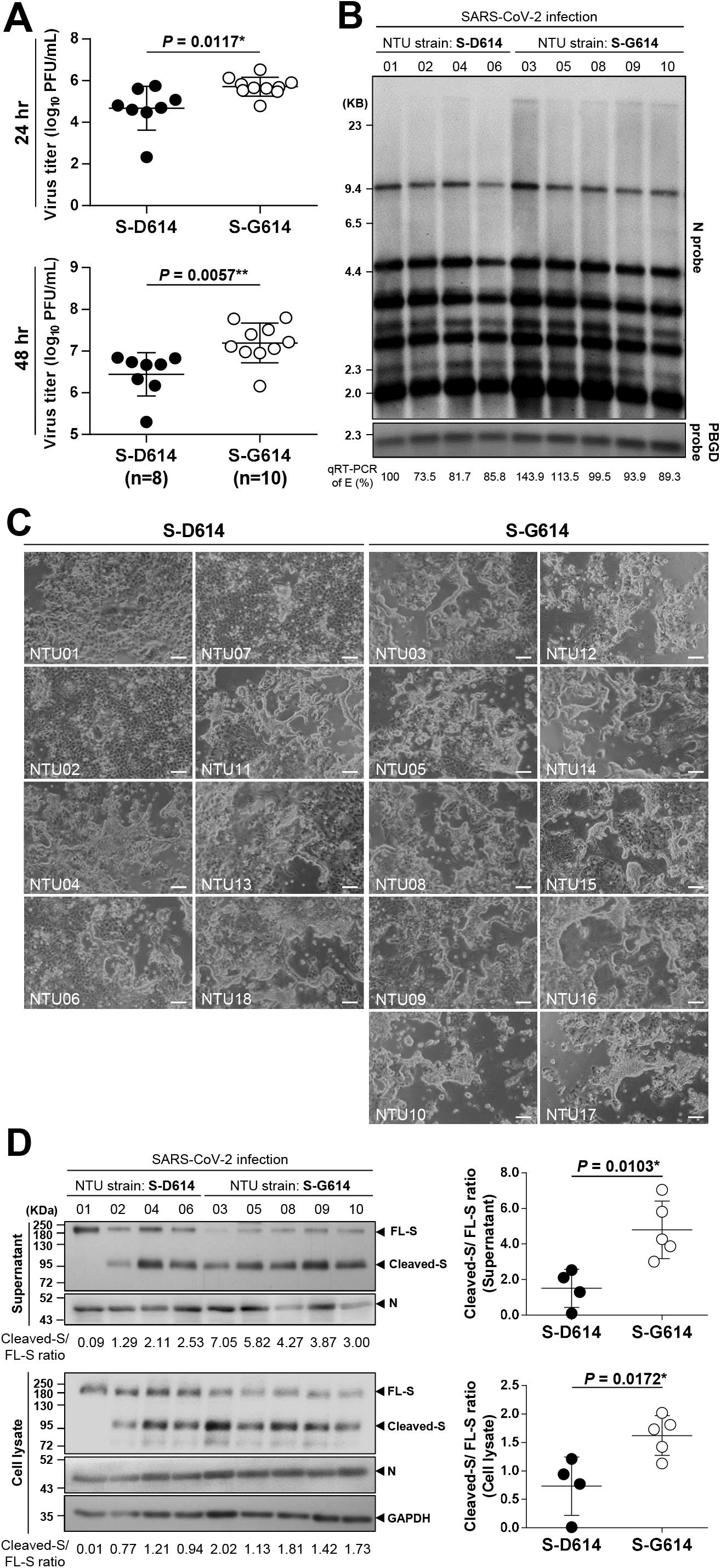
Virus production and cytopathic syncytium formation were higher for S-G614 containing SARS-CoV-2 than for S-D614 containing SARS-CoV-2, associated with an increase in cleavage of the S protein. **(A)** Comparison of the virus titers (plaque-forming units (PFU)/mL) (MOI=0.01) in the supernatants of Vero E6 cells infected with SARS-CoV-2 strains NTU01-NTU18 (8 strains expressing wild-type S-D614 and 10 strains expressing mutant S-G614) at 24 hr (upper panel) and 48 hr (lower panel) postinfection. Data are represented as the mean ± SD (*P* < 0.05*; *P* < 0.01**). **(B)** The representative results of northern blot analysis of viral RNA isolated from Vero E6 cells infected with S-D614 or S-G614 containing viruses at 48 hr postinfection. Viral RNA was quantified by qRT-PCR targeting the E gene as indicated below the northern blot results. **(C)** Microscopic observation of syncytia in Vero E6 cells infected with the SARS-CoV-2 strains NTU01-NTU18 (MOI=0.01), which expressed either the S-D614 protein (left panel) or S-G614 protein (right panel), at 48 hr postinfection. Scale bars: 100 μm. **(D)** Left panel: representative immunoblot of S protein extracted from the supernatants and cell lysates of S-D614-cotnaining (lanes 1-4) or S-G614-containing (lanes 5-9) virus-infected Vero E6 cells at 48 hr postinfection (MOI=0.01). Full length (FL) S proteins, cleaved S proteins and nucleocapsid (N) proteins are marked as indicated. The ratio of cleaved S to FL S protein for each SARS-CoV-2 strain indicated below the immunoblot results. The ratios in the S-D614- and S-G614-containing viruses were compared and are presented as the mean ± SD in the right panel (*P* < 0.05*).

### S-G614-containing viral isolates showed increased cleavage of the S protein than S-D614-containing viral isolates

As shown in our previous study, cleavage of the S protein at the S1/S2 boundary by furin/PCs is critical for S-mediated syncytium formation (Cheng et al., 2020). We thus compared the patterns of S protein cleavage at this site for viruses containing either the S-G614 or S-D614 protein. Viruses in the supernatants of cells infected with either group of viruses were harvested for immunoblot analysis. Interestingly, the results showed significantly increased cleavage of the S protein into the S1 and S2 fragments in S-G614-containing viruses than in S-D614-containing viruses, and the cleaved S/full-length (FL) S ratio was shown to be 4.8 ± 0.7 versus 1.5 ± 0.5, respectively (*P*=0.0103) (Figure 1D, upper panel). Consistently, a significant difference in cleavage of S between lysates from cells infected with the two groups of viruses was also demonstrated by immunoblotting, with the cleaved S/FL S ratio found to be 1.6 ± 0.2 versus 0.7 ± 0.3, respectively (*P*=0.0172) (Figure 1D, lower panel). Therefore, these results suggested the function of the D614G substitution in enhancing cleavage of the S protein.

### The syncytial phenotype and S cleavage were increased in S-G614-expressing cells than in S-D614-expressing cells

Genetic heterogeneity beyond the D614G substitution among different SARS-CoV-2 isolates might cause confusion regarding our observation. Therefore, we tested our hypothesis in cultured cells expressing only the S protein. Codon-optimized expression plasmids for wild-type S-D614 and the single substitution mutant (S-G614) were individually transfected into Vero E6 cells, which were harvested at 24 hr posttransfection for analysis. We first examined the syncytial phenotype via the observation of fused cells containing multiple nuclei visible under light microscopy. Compared with control cells transfected with only the vector, cells expressing the S-D614 protein did exhibit a moderate syncytial phenotype, which was increased in cells expressing the mutant S-G614 protein (Figure 2A). Immunoblot analysis further revealed increased cleavage of the S-G614 proteins relative to the S-D614 protein and a higher cleaved S/FL S ratio in the lysate of S-G614-expressing cells compared to S-D614-expressing cells, not only at 24 hr posttransfection but also at 16 and 20 hr posttransfection (0.13 vs 0.45, 0.29 vs 0.66, and 0.41 vs 0.93 at 16, 20 and 24 hr posttransfection, respectively) (Figure 2B). These results are consistent with the findings from the virus infection system shown in Figure 1D.

**Figure 2.**
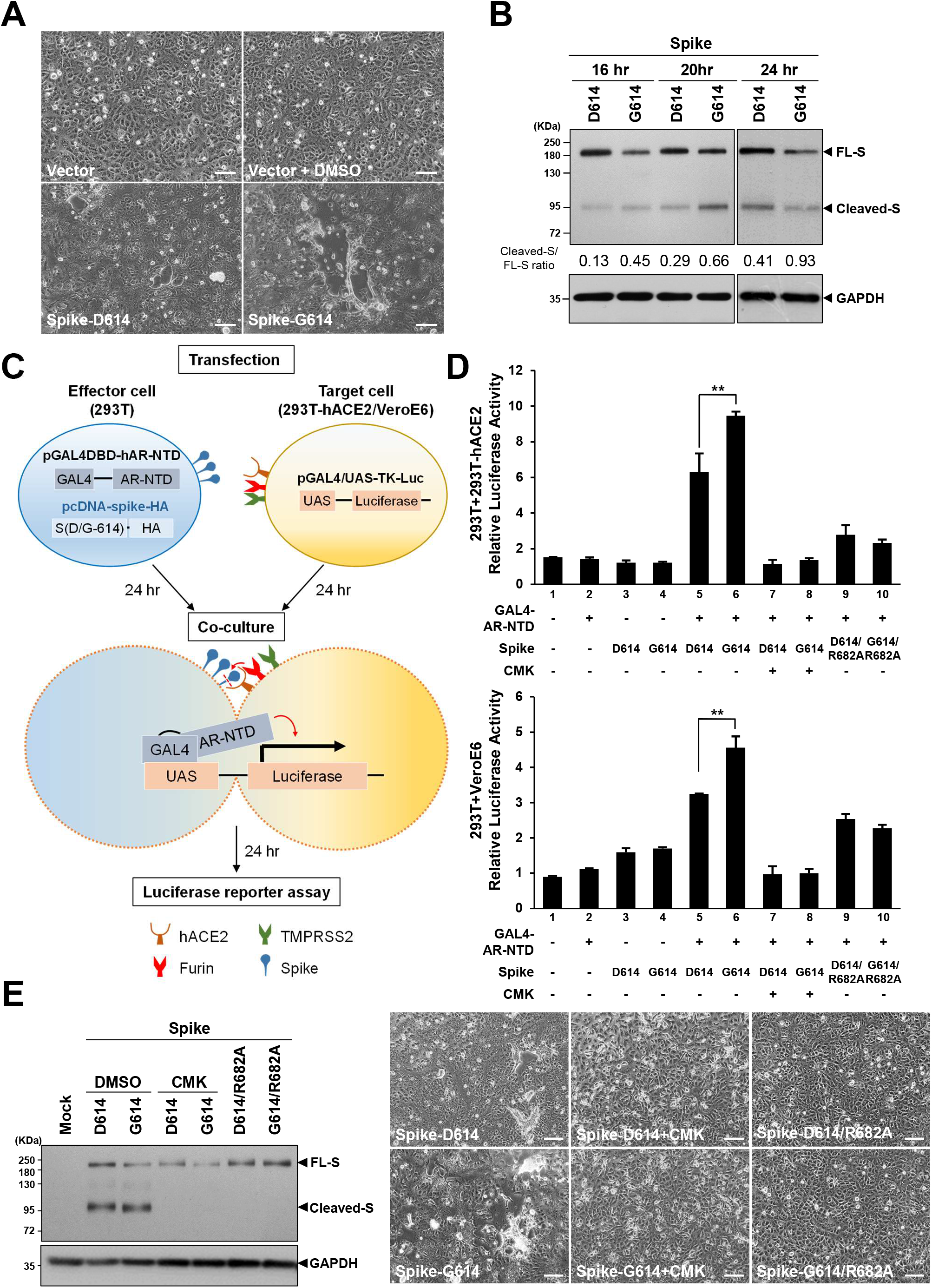
The fusion activity of the SARS-CoV-2 S protein was higher in S-G614-expressing cells than in S-D614-expressing cells and dependent on furin/PC mediated cleavage of S protein. **(A)** Microscopic observation of syncytia in Vero E6 cells expressing the S-D614 or S-G614 protein. Scale bars: 100 μm. **(B)** Immunoblot analysis of lysates from Vero E6 cells transfected with expression constructs for S-D614 or S-G614 harvested at 16-24 hr posttransfection. The immunoblot was probed with anti-S Ab, and the full-length (FL) and cleaved S proteins are marked as indicated; GAPDH was included as a loading control. The ratio of cleaved S versus FL S is indicated below the immunoblot results. **(C)** Schematic illustration of the one-hybrid luciferase reporter assay designed to quantitatively evaluate the fusion activity induced by SARS-CoV-2 S protein. Effector 293T cells were cotransfected with an expression plasmid for S-D614 or S-G614 and the pGAL4DBD-hAR-NTD plasmid. The target 293T-hACE2 or Vero E6 cells were transfected with pGAL4/UAS-TK-Luc. At 24 hr posttransfection, the effector and target cells were cocultured for 24 hr and harvested to assay the luciferase activity. **(D)** Representative results of the luciferase reporter assay showing luciferase activity induced by S-D614 or S-G614 protein (upper panel, in 293T-hACE2 target cells; lower panel, in Vero E6 target cells). The luciferase activity in cells expressing the SARS-CoV-2 S-D614/S-G614 protein or D614/R682A or G614/R682A protein with or without treatment with a furin/PC inhibitor (CMK, 50 μM) was assessed as indicated. The results were derived from three independent experiments and are shown as the mean ±SD (*P* < 0.01**). **(E)** The effect of treatment with CMK (50 μM) and the R682A mutation of S on syncytium formation induced by S-D614 or S-G614 protein. Left panel, immunoblot analysis of the lysates from Vero E6 cells transfected with the indicated plasmids with or without CMK treatment. Right panel, microscopic observation of cell morphology in Vero E6 cells transfected with the indicated plasmids with or without CMK treatment. Scale bars: 100 μm

### The S mediated syncytium was increased in S-G614-expressing cells than in S-D614-expressing cells, dependent on furin-mediated S cleavage

To confirm the effect of D614G substitution on S and ACE2 binding-mediated cell syncytium formation, we established a luciferase-based reporter assay to quantitatively compare syncytium induction by the S-G614 and S-D614 proteins (schematically illustrated in Figure 2C). As documented, interaction between S protein and ACE2 at the membrane of infected cells and adjacent cells primes the cleavage of S by host proteases to release the fusion peptide essential for syncytium formation (Bertram et al., 2011). Therefore, in our experimental design, we selected ACE2-null 293T cells transfected with expression plasmid for either S-D614 or S-G614 as S(+) and ACE2(-) effector cells. Two ACE2(+) target cell lines were selected: 293T cells stably transfected with human ACE2 (293T-hACE2) and Vero E6 cells (with endogenous ACE2 expression). Neither effector cells nor target cells alone developed syncytia due to the lack of the binding partner (ACE2 or the S protein). To quantitatively examine the fusion of effector and target cells, we developed a one hybrid luciferase reporter assay. The pGAL4DBD-hAR-NTD plasmid was co-transfected with an individual S expression plasmid into the effector cells; the pGAL4/UAS-TK-Luc plasmid was transfected into the target cells. The effector and target cells were then co-cultured and harvested for reporter activity analysis. The expression of pGAL4/UAS-TK-Luc in the target cells was activated only upon formation of syncytia consisting of effector and target cells (mediated by interaction of S and ACE2), which is driven by the transcriptional activator encoded by the pGAL4DBD-hAR-NTD plasmid in the effector cells.

As expected, the expression of pGAL4DBD-hAR-NTD or S alone in the effector cells did not activate luciferase expression in the target cells (Figure 2D, lanes 2-4), which was elevated only when pGAL4DBD-hAR-NTD and the S protein were coexpressed in the effector cells. Expression of S-G614 significantly increased luciferase activity compared to that upon the expression of S-D614 in the effector cells (Figure 2D, lane 5 vs 6). The results from the reporter assay thus provide quantitative evidence supporting increased syncytium induction by the mutant S-G614 protein compared with the S-D614 protein, consistent with the findings from light microscopy observation.

To further examine whether the enhanced syncytium formation by S-G614 expression was mediated through cleavage by furin/PC protease(s), we treated S-expressing effector cells with the furin/PC inhibitor decanoyl-RVKR-chloromethylketone (CMK). Immunoblot analysis demonstrated that cleavage of the S-D614 or S-G614 protein was completely blocked by CMK treatment (Figure 2E, left panel). The syncytial phenotype in the S-D614- and S-G614-expressing cells was also significantly decreased by CMK treatment (Figure 2E, right panel). The difference in luciferase activity induced by the S-D614 and S-G614 proteins was diminished by CMK treatment (Figure 2D, lane 7 vs 8), suggesting that the enhanced syncytium formation induced by the S-G614 protein is dependent on the presence of active furin/PCs in cells. Consistent with this finding, the difference in the effects of S-D614 and S-G614 on reporter activity was abolished when the furin substrate site was mutated by introduction of the R682A substitution (Figure 2D, lane 9 vs 10); the effects of the substitution on cleavage of the S protein and syncytium formation are shown in Figure 2E. Altogether, these results suggest that the putative function of the D614G mutation in the S protein of SARS-CoV-2 is dependent on enhanced cleavage at the furin substrate motif at S1/S2 boundary, which contributes to an increased membrane fusion activity.

## DISCUSSION

Both virus entrance and syncytium formation during SARS-CoV-2 infection are mediated through membrane fusion between the virus and cell or between cells. As shown in our recent study, cleavage of the S protein by furin/PCs is critical for the membrane fusion of SARS-CoV-2, after the binding of S protein with ACE2 receptor (Cheng et al., 2020). The current study further revealed that the D614G substitution in S contributes to increasing accessibility to the polybasic RRAR motif for cleavage by furin/PCs, leading to the increased infectivity and syncytium formation of S-G614-containing SARS-CoV-2 viruses. In fact, syncytium formation has also been documented to confer advantages in terms of infectivity and transmission in many coronaviruses (CoVs) (Frana et al., 1985; Nakagaki et al., 2005; Park et al., 2016; Yamada and Liu, 2009). Direct cell-to-cell spread of CoVs is more efficient than their cell-free spread; the syncytium also allows viruses to evade innate immune surveillance (Sattentau, 2008, 2011). Moreover, an unique structure generated by cleavage of S at furin substrate site was identified to be critical for viral entry and infectivity, via interacting with cell surface Neurophilin-1 (NRP1) (Cantuti-Castelvetri et al., 2020; Daly et al., 2020). Therefore, the effect of the D614G substitution in increasing cleavage of the furin substrate motif could benefit viral infectivity and transmission and provides a mechanism for the significant expansion of S-G614-containing SARS-CoV-2 worldwide.

Through cryo-EM analysis, Yurkovetskiy *et al.* reported that the D614G mutation might introduce a conformational change in the spike protein (Yurkovetskiy et al., 2020), implying an improved ability to bind to ACE2 receptor, a key for subsequent membrane fusion. More recently, Hou *et al.* showed that the D614G variant SARS-CoV-2 replicates more efficiently in primary human proximal airway epithelial cells than the wild-type virus. They also provided *in vivo* evidence showing that the D614G variant exhibited significantly faster droplet transmission between hamsters than the wild-type virus at early stage after infection (Hou et al., 2020a; Plante et al., 2020; Yurkovetskiy et al., 2020). Our study here provides mechanistic explanation in support of the notion that the D614G substitution can increase virus transmission through enhanced membrane fusion mediated by increased cleavage of spike by furin/PC proteases. Though this mechanism was delineated in assays of cells expressing only the S protein, it has been demonstrated in virus-infected cells, as some S-G614-containing viruses did show more evident cleavage of the S protein (Figure 1D). Despite this, the D614G mutation was noted to be associated with mutations in the viral nsp3 and an ORF1b protein variant (P314L) (Bhattacharyya et al., 2020; Mercatelli and Giorgi, 2020). The coexistence of D614G and P314L may suggest additional mechanisms for the selection advantage of viruses expressing S-G614 in terms of viral infectivity. In addition, the effect of D614G might be confounded by other genetic variations among different virus isolates.

Currently, S-G614-containing SARS-CoV-2 has expanded and is the dominant strain worldwide. As the enhancement of cleavage by furin/PCs is critical for its transmission, blockade of this mechanism by targeting furin/PC protease activity for inhibition might become a potential antiviral strategy to block transmission of this strain. As shown in our recent study, two furin/PC inhibitors, CMK and naphthofluorescein, can abolish S cleavage, virus production and pathogenic syncytium formation of an S-G614-containing SARS-CoV-2 isolate (NTU03) (Cheng et al., 2020). Such inhibitors thus become leads for further antiviral development for the prevention and treatment of S-G614-containing SARS-CoV-2 infection.

## ACKNOWLEDGMENTS

This study was supported by grants from the Ministry of Science and Technology, Taiwan (MOST109-2327-B-002-009, M0ST109-2634-F-002-043, M0ST109-3114-Y-001-001) and the “Center of Precision Medicine” from The Featured Areas Research Center Program within the framework of the Higher Education Sprout Project by the Ministry of Education (MOE) in Taiwan. We would like to acknowledge the service provided by the Biosafety Level-3 Laboratory of the First Core Laboratory, National Taiwan University College of Medicine.

## AUTHOR CONTRIBUTION

Y.-W.C., S.-H.Y., and S.-Y.C. designed research; Y.-W.C. performed SARS-CoV-2 S transfection experiments; T.-L.C. performed viral infection experiments. H.-C.K. and Y.-M.T. contributed to the viral infection experiments, plaque assay and virus titer determination; Y.-W.C. cloned SARS-CoV-2 S-D614G-HA, S-R682A-HA, and S-D614G/R682A-HA. Y.-W.C. and C.-L.H. performed western and northern blotting; S.-H.W. contributed to northern blotting; C.-L.L. performed qRT-PCR. Y.-W.C., C.-L.L., S.-H.Y., and S.-Y.C. analyzed data; Y.-W.C., S.-Y.C., and S.-H.Y. drafted the manuscript; H.-Y.W., P.-J.C. critical revision of the manuscript.

## DECLARATION OF INTERESTS

The authors declare no competing interests.

## STAR METHODS

### RESOURCE AVAILABILITY

#### Lead Contact

Further information and requests for resources and reagents should be directed to and will be fulfilled by the Lead Contact, Shiou-Hwei Yeh

#### Materials Availability

All materials and reagents will be made available upon instalment of a material transfer agreement (MTA).

#### Data and Code Availability

The original sequencing datasets for hCoV-19/Taiwan/NTU01/2020 to hCoV-19/Taiwan/NTU18/2020 can be found on the GISAID under Accession ID listed in Table S1.

### EXPERIMENTAL MODEL AND SUBJECT DETAILS

#### Viruses

Sputum specimens from SARS-CoV-2-infected patients were kept in viral transport medium. Virus isolated from the specimens was propagated in Vero E6 cells in Dulbecco’s modified Eagle’s medium (DMEM) supplemented with 2 μg/mL tosylsulfonyl phenylalanyl chloromethyl ketone-trypsin. The 18 virus isolates used in the current study were hCoV-19/Taiwan/NTU01/2020 to hCoV-19/Taiwan/NTU18/2020. The sequencing data have been deposited in the GISAID, and the accession codes are listed in Table S1.

#### Plaque assay

The plaque assay was performed as previously described with minor modifications (Su et al., 2008). In brief, Vero E6 cells (2 x 10^5^ cells/well) were seeded in 24-well tissue culture plates and maintained in DMEM supplemented with 10% FBS and antibiotics. After 24 hr incubation, SARS-CoV-2 virus was treated to the cell monolayer for 1 hr at 37°C. After removed the virus and washed the cell monolayer once with PBS, maintained the cells with medium containing 1% methylcellulose and incubated for 5-7 days. Subsequently, cells fixed with 10% formaldehyde overnight and stained with 0.7% crystal violet in order to count the plaques. The virus titer is from the mean of three independent experiments.

#### Plasmid construction

Humanized pUC57-2019-nCoV-S was a kind gift from Dr. Che Ma at the Institute of Genomics Research Center, Academia Sinica, Taiwan. The spike sequence was cloned into the pcDNA3.0-HA vector with the addition of an HA tag at the spike protein C-terminus via NheI and XbaI sites. A QuikChange II site-directed mutagenesis kit (Agilent) was used to generate the mutant D614G-spike or R682A-spike construct. The primer set for D614G-spike was S-D614G-F: 5’-CTCGGTACAATTCACGCCCTGATACAGCACGGC-3’ and S-D614G-R: 5’-GCCGTGCTGTATCAGGGCGTGAATTGTACCGAG-3’. The primer set for R682A-spike was 5’-ACGCTCCGGGCTCTTGCGGGAGAGTTTGTCTG-3’ and 5’-CAGACAAACTCTCCCGCAAGAGCCCGGAGCGT-3’.

#### Cell culture experiments

293T cells stably expressing human ACE2 (293T-ACE2) were kindly provided from professor Mi-Hua Tao. The 293T, 293T-hACE2 and VeroE6 cells were maintained and grown at 37°C in Dulbecco’s Modified Eagle’s Medium (DMEM, Biological Industries) containing 10% fetal bovine serum (FBS) (HyClone, GE Healthcare Life Sciences) in a 5% CO_2_ incubator. The spike-D614 or spike-G614 plasmid was transfected into VeroE6 cells with Lipofectamine™ 2000 (Thermo Fisher Scientific). The furin inhibitor, CMK (10 mM in DMSO, final concentration is 50 μM in 0.5% DMSO (v/v)) (Tocris Bioscience), was added to the medium at the indicated concentrations 2 hr post transfection. Cells were harvested 24 hr posttransfection for subsequent western blot.

#### Cell-cell fusion assay

A quantitative GAL4-based mammalian one-hybrid assay was established to assess cell-cell fusion activity. This reporter assay contains two plasmids (kindly provided by Dr. Hsiu-Ming Shih at the Institute of Biomedical Sciences, Academia Sinica, Taiwan). One is the reporter construct, pGAL4/UAS-TK-Luc, which is the firefly luciferase under the control of the GAL4 response element (UAS) and a thymidine kinase (TK) promoter. The other is the transcriptional activator construct, pGAL4DBD-hAR-NTD, which consists of a.a. 1-560 of the AR transcriptional activation domain with the GAL4 DNA-binding domain (DBD) fused at its N-terminus. The protein encoded by pGAL4DBD-hAR-NTD can bind to the UAS GAL4 response element of pGAL4/UAS-TK-Luc reporter and activate the transcription of the luciferase reporter gene by the transcriptional activator of AR-NTD.

To detect cell-cell fusion activity, 293T-hACE2 and Vero E6 (which endogenously express ACE2) cells transfected with pGAL4/UAS-TK-Luc were prepared as target cells; 293T cells expressing the SARS-CoV-2 spike protein and pGAL4DBD-hAR-NTD were prepared as effector cells. In brief, 3×10^5^ 293T cells were seeded in 12-well plates, and Vero E6 and 293T-hACE2 cells were seeded in 24-well plates (2×10^5^/well) overnight. After 24 hr, the 293T cells were cotransfected with pGAL4DBD-hAR-NTD and D614-spike or G614-spike plasmid. Vero E6 cells and 293T-hACE2 cells were transfected with pGAL4/UAS-TK-Luc and pCMV-Renilla. Twenty-four hours post transfection, 293T cells expressing the GAL4DBD-hAR-NTD protein and D614-spike or G614-spike protein were suspended in trypsin, and 1×10^5^ cells were seeded on Vero E6 or 293T-hACE2 cells expressing the GAL4/UAS-TK-Luc protein. The cells were co-cultured for 24 hr and harvested with passive lysis buffer (PLB) for the dual-luciferase assay following the manufacture’s instruction (Promega).

#### Western blot analysis

Western blotting was performed as previously described (Wu et al., 2009). In brief, cell lysates were extracted by 1× RIPA buffer (Merck Millipore) containing 1× proteinase inhibitor (Merck Millipore) and 1× phosphatase inhibitor (Calbiochem). Equal amounts of protein samples were electrophoretically separated by 10% sodium dodecyl sulfate polyacrylamide gel electrophoresis (SDS-PAGE) and transferred to polyvinylidene difluoride (PVDF) membranes. The membranes were probed with the indicated primary antibodies at 4°C overnight and then reacted with a secondary antibody. Antigen-antibody complexes were visualized using Western Lightning Plus-ECL (PerkinElmer). The antibodies used for western blot analysis were as follows: rabbit anti-SCoV/SARS-CoV-2 nucleocapsid (generated by our laboratory), mouse anti-SARS-CoV/SARS-CoV-2 (COVID-19) spike [1A9] (Genetex, GTX632604), rabbit anti-GAPDH (Genetex, GTX100118), horseradish peroxidase-conjugated mouse IgG (Genetex, GTX213111-01) and rabbit IgG (Genetex, GTX213110-01). The quantitative result of cleaved to full-length spike ratio in immunoblot was analyzed by VisionWorks Life Science Image Analysis software (UVP, Upland, CA USA)

#### RNA extraction and northern blot analysis

RNA was extracted by NucleoSpin RNA Kit (Macherey-Nagel) according to the instruction. Northern blotting was performed as previously described (Wu et al., 2014). In brief, 0.2 μg of RNA was denatured and separated by an 0.8% agarose/formaldehyde gel at 70 volts for 5 hr. The agarose gel was soaked in 50 mM NaOH for 50 min to break the large RNA fragment. Subsequently, gel washed with 100 mM Tris-HCl (pH 7.5) for 30 min, and incubated in 20× SSC buffer for 20 min and then capillary-transferred to a positively charged Hybond-N nylon membrane (Amersham Biosciences) overnight. RNA was immobilized by UV crosslinking (1800 x 100 μJ/cm) and hybridized at 50°C overnight with digoxigenin (DIG)-labeled probes generated with a PCR DIG probe synthesis kit (Roche Diagnostics). The primers used to synthesize the DIG-labeled nCoV19-N cDNA probe were 5’-AAGCTGGACTTCCCTATGGTGC-3’ and 5’-CCTTGGGTTTGTTCTGGACCACG-3’. The probes of porphobilinogen deaminase (PBGD) used as the internal control in northern blot were 5’-GGTGACCAGCACACTTTGGG-3’ and 5’-AGCCGGGTGTTGAGGTTTCC-3’.

#### Quantitative reverse transcription polymerase chain reaction (qRT-PCR)

The qRT-PCR was conducted following the protocol as described previously (Cheng et al., 2020). In brief, RNA extracted from SARS-CoV-2 infected VeroE6 cells was reverse transcribed using SuperScript III Reverse Transcriptase System (Thermo Fisher Scientific). Quantitative PCR targeting to E gene was performed using FastStart DNA SYBR Green on LightCycler 1.5 (Roche Diagnostics), with primer set of 5’-ACAGGTACGTTAATAGTTAATAGCGT-3’ and 5’-ATATTGCAGCAGTACGCACACA-3’. The RNA level of E gene in the cells was determined in relative to the internal control of cellular PBGD gene, with primer set of 5’-GCATCGCTGAAAGGGCCTTCC-3’ and 5’-TCATCCTCAGGGCCATCTTCATGC-3’.

#### Statistical analysis

The virus titers quantified by plaque assays in triplicate are shown as the mean ± SD. Results from the reporter assay are shown as data representative of three independent experiments and presented as the mean ± SD. Differences in data from the virus titer, qRT-PCR and reporter assay between each indicated paired sample groups were evaluated by Student’s t-test. A P value of 0.05 or lower was used to indicate statistical significance (*, *P* < 0.05; **, *P* < 0.01; ***, *P* < 0.001).

## Notes

### Competing Interest Statement

The authors have declared no competing interest.

## REFERENCES

Bertram, S., Glowacka, I., Muller, M.A., Lavender, H., Gnirss, K., Nehlmeier, I., Niemeyer, D., He, Y., Simmons, G., Drosten, C., et al. (2011). Cleavage and activation of the severe acute respiratory syndrome coronavirus spike protein by human airway trypsin-like protease. J Virol 85, 13363–13372.

Bhattacharyya, C., Das, C., Ghosh, A., Singh, A.K., Mukherjee, S., Majumder, P.P., Basu, A., and Biswas, N.K. (2020). Global spread of SARS-CoV-2 subtype with spike protein mutation D614G is shaped by human genomic variations that regulate expression of TMPRSS2 and MX1 genes. bioRxiv, 2020.05.04.075911.

Cantuti-Castelvetri, L., Ojha, R., Pedro, L.D., Djannatian, M., Franz, J., Kuivanen, S., van der Meer, F., Kallio, K., Kaya, T., Anastasina, M., et al. (2020). Neuropilin-1 facilitates SARS-CoV-2 cell entry and infectivity. Science.

Chellapandi, P., and Saranya, S. (2020). Genomics insights of SARS-CoV-2 (COVID-19) into target-based drug discovery. Med Chem Res 29, 1777–1791.

Cheng, Y.W., Chao, T.L., Li, C.L., Chiu, M.F., Kao, H.C., Wang, S.H., Pang, Y.H., Lin, C.H., Tsai, Y.M.’ Lee, W.H., et al. (2020). Furin inhibitors block SARS-CoV-2 spike protein cleavage to suppress virus production and cytopathic effects. Cell Rep 33, 108254.

Chitranshi, N., Gupta, V.K., Rajput, R., Godinez, A., Pushpitha, K., Shen, T., Mirzaei, M., You, Y.Y., Basavarajappa, D., Gupta, V., et al. (2020). Evolving geographic diversity in SARS-CoV2 and in silico analysis of replicating enzyme 3CL(pro)targeting repurposed drug candidates. J Transl Med 18, 10.1186/s12967-12020-02448-z.

Daly, J.L., Simonetti, B., Klein, K., Chen, K.E., Williamson, M.K., Anton-Plagaro, C., Shoemark, D.K., Simon-Gracia, L., Bauer, M., Hollandi, R., et al. (2020). Neuropilin-1 is a host factor for SARS-CoV-2 infection. Science.

Daniloski, Z., Guo, X., and Sanjana, N.E. (2020). The D614G mutation in SARS-CoV-2 Spike increases transduction of multiple human cell types. bioRxiv, 2020.06.14.151357.

Frana, M.F., Behnke, J.N., Sturman, L.S., and Holmes, K.V. (1985). Proteolytic cleavage of the E2-glycoprotein of murine coronavirus - host-dependent differences in proteolytic cleavage and cell-fusion. Journal of Virology 56, 912–920.

Hoffmann, M., Kleine-Weber, H., and Pohlmann, S. (2020). A multibasic cleavage site in the spike protein of SARS-CoV-2 is essential for infection of human lung cells. Mol Cell 78, 1–6.

Hou, Y.J., Chiba, S., Halfmann, P., Ehre, C., Kuroda, M., Dinnon, K.H., Leist, S.R., Schäfer, A., Nakajima, N., Takahashi, K., et al. (2020a). SARS-CoV-2 D614G variant exhibits enhanced replication *ex vivo* and earlier transmission *in vivo*. bioRxiv, 2020.09.28.317685.

Hou, Y.J., Okuda, K., Edwards, C.E., Martinez, D.R., Asakura, T., Dinnon, K.H., 3rd, Kato, T., Lee, R.E., Yount, B.L., Mascenik, T.M., et al. (2020b). SARS-CoV-2 reverse genetics reveals a variable infection gradient in the respiratory tract. Cell 182, 429–446 e414.

Hu, J., He, C.-L., Gao, Q.-Z., Zhang, G.-J., Cao, X.-X., Long, Q.-X., Deng, H.-J., Huang, L.-Y., Chen, J., Wang, K., et al. (2020). D614G mutation of SARS-CoV-2 spike protein enhances viral infectivity. bioRxiv, 2020.06.20.161323.

Korber, B., Fischer, W.M., Gnanakaran, S., Yoon, H., Theiler, J., Abfalterer, W., Hengartner, N., Giorgi, E.E., Bhattacharya, T., Foley, B., et al. (2020). Tracking changes in SARS-CoV-2 spike: Evidence that D614G increases infectivity of the COVID-19 virus. Cell 182, 812–827.

Mercatelli, D., and Giorgi, F.M. (2020). Geographic and genomic distribution of SARS-CoV-2 mutations. Front Microbiol 11.

Nakagaki, K., Nakagaki, K., and Taguchi, F. (2005). Receptor-independent spread of a highly neurotropic murine coronavirus JHMV strain from initially infected microglial cells in mixed neural cultures. J Virol 79, 6102–6110.

Ou, X., Zheng, W., Shan, Y., Mu, Z., Dominguez, S.R., Holmes, K.V., and Qian, Z. (2016). Identification of the fusion peptide-containing region in betacoronavirus spike glycoproteins. J Virol 90, 5586–5600.

Pachetti, M., Marini, B., Benedetti, F., Giudici, F., Mauro, E., Storici, P., Masciovecchio, C., Angeletti, S., Ciccozzi, M., Gallo, R.C., et al. (2020). Emerging SARS-CoV-2 mutation hot spots include a novel RNA-dependent-RNA polymerase variant. J Transl Med 18, 179.

Park, J.E., Li, K., Barlan, A., Fehr, A.R., Perlman, S., McCray, P.B., Jr., and Gallagher, T. (2016). Proteolytic processing of Middle East respiratory syndrome coronavirus spikes expands virus tropism. Proc Natl Acad Sci U S A 113, 12262–12267.

Plante, J.A., Liu, Y., Liu, J., Xia, H., Johnson, B.A., Lokugamage, K.G., Zhang, X., Muruato, A.E., Zou, J., Fontes-Garfias, C.R., et al. (2020). Spike mutation D614G alters SARS-CoV-2 fitness and neutralization susceptibility. bioRxiv, 2020.09.01.278689.

Sanjuan, R., and Domingo-Calap, P. (2016). Mechanisms of viral mutation. Cell Mol Life Sci 73, 4433–4448.

Sattentau, Q.J. (2008). Avoiding the void: cell-to-cell spread of human viruses. Nat Rev Microbiol 6, 815–826.

Sattentau, Q.J. (2011). The direct passage of animal viruses between cells. Curr Opin Virol 1, 396–402.

Su, C.T., Hsu, J.T., Hsieh, H.P., Lin, P.H., Chen, T.C., Kao, C.L., Lee, C.N., and Chang, S.Y. (2008). Anti-HSV activity of digitoxin and its possible mechanisms. Antiviral Res 79, 62–70.

Weissman, D., Alameh, M.-G., de Silva, T., Collini, P., Hornsby, H., Brown, R., LaBranche, C.C., Edwards, R.J., Sutherland, L., Santra, S., et al. (2020). D614G spike mutation increases SARS CoV-2 susceptibility to neutralization. medRxiv, 2020.07.22.20159905.

Wu, C.H., Chen, P.J., and Yeh, S.H. (2014). Nucleocapsid phosphorylation and RNA helicase DDX1 recruitment enables coronavirus transition from discontinuous to continuous transcription. Cell Host Microbe 16, 462–472.

Wu, C.H., Yeh, S.H., Tsay, Y.G., Shieh, Y.H., Kao, C.L., Chen, Y.S., Wang, S.H., Kuo, T.J., Chen, D.S., and Chen, P.J. (2009). Glycogen synthase kinase-3 regulates the phosphorylation of severe acute respiratory syndrome coronavirus nucleocapsid protein and viral replication. Journal of Biological Chemistry 284, 5229–5239.

Yamada, Y., and Liu, D.X. (2009). Proteolytic activation of the spike protein at a novel RRRR/S motif is implicated in furin-dependent entry, syncytium formation, and infectivity of coronavirus infectious bronchitis virus in cultured cells. Journal of Virology 83, 8744–8758.

Yin, C. (2020). Genotyping coronavirus SARS-CoV-2: methods and implications. Genomics 112, 3588–3596.

Young, B.E., Fong, S.W., Chan, Y.H., Mak, T.M., Ang, L.W., Anderson, D.E., Lee, C.Y., Amrun, S.N., Lee, B., Goh, Y.S., et al. (2020). Effects of a major deletion in the SARS-CoV-2 genome on the severity of infection and the inflammatory response: an observational cohort study. Lancet 396, 603–611.

Yurkovetskiy, L., Wang, X., Pascal, K.E., Tomkins-Tinch, C., Nyalile, T.P., Wang, Y., Baum, A., Diehl, W.E., Dauphin, A., Carbone, C., et al. (2020). Structural and functional analysis of the D614G SARS-CoV-2 spike protein variant. Cell 183.

