## Supplemental Table 1 for "D614G Substitution of SARS-CoV-2 Spike Protein Increases Syncytium Formation and Viral Transmission via Enhanced Furin-mediated Spike Cleavage"

### Supplemental Information

**Table S1: SARS-CoV-2 genome (GISAID ID)**

| Virus name | Accession ID | Location |
|---|---|---|
| hCoV-19/Taiwan/NTU01/2020 | EPI_ISL_408489 | Asia / Taiwan / Taipei |
| hCoV-19/Taiwan/NTU02/2020 | EPI_ISL_410218 | Asia / Taiwan / Taipei |
| hCoV-19/Taiwan/NTU03/2020 | EPI_ISL_413592 | Asia / Taiwan / Taipei |
| hCoV-19/Taiwan/NTU04/2020 | EPI_ISL_422407 | Asia / Taiwan / Taipei |
| hCoV-19/Taiwan/NTU05/2020 | EPI_ISL_422408 | Asia / Taiwan / Taipei |
| hCoV-19/Taiwan/NTU06/2020 | EPI_ISL_422409 | Asia / Taiwan / Taipei |
| hCoV-19/Taiwan/NTU07/2020 | EPI_ISL_422410 | Asia / Taiwan / Taipei |
| hCoV-19/Taiwan/NTU08/2020 | EPI_ISL_422411 | Asia / Taiwan / Taipei |
| hCoV-19/Taiwan/NTU09/2020 | EPI_ISL_422412 | Asia / Taiwan / Taipei |
| hCoV-19/Taiwan/NTU10/2020 | EPI_ISL_447614 | Asia / Taiwan / Taipei |
| hCoV-19/Taiwan/NTU11/2020 | EPI_ISL_422413 | Asia / Taiwan / Taipei |
| hCoV-19/Taiwan/NTU12/2020 | EPI_ISL_422414 | Asia / Taiwan / Taipei |
| hCoV-19/Taiwan/NTU13/2020 | EPI_ISL_422415 | Asia / Taiwan / Taipei |
| hCoV-19/Taiwan/NTU14/2020 | EPI_ISL_422416 | Asia / Taiwan / Taipei |
| hCoV-19/Taiwan/NTU15/2020 | EPI_ISL_422417 | Asia / Taiwan / Taipei |
| hCoV-19/Taiwan/NTU16/2020 | EPI_ISL_422418 | Asia / Taiwan / Taipei |
| hCoV-19/Taiwan/NTU17/2020 | EPI_ISL_422419 | Asia / Taiwan / Taipei |
| hCoV-19/Taiwan/NTU18/2020 | EPI_ISL_447615 | Asia / Taiwan / Taipei |
